# Hippocampal Spike-Timing Correlations Lead to Hexagonal Grid Fields

**DOI:** 10.1101/153379

**Authors:** Mauro M Monsalve-Mercado, Christian Leibold

## Abstract

Space is represented in the mammalian brain by the activity of hippocampal place cells as well as in their spike-timing correlations. Here we propose a theory how this temporal code is transformed to spatial firing rate patterns via spike-timing-dependent synaptic plasticity. The resulting dynamics of synaptic weights resembles well-known pattern formation models in which a lateral inhibition mechanism gives rise to a Turing instability. We identify parameter regimes in which hexagonal firing patterns develop as they have been found in medial entorhinal cortex.

---

The spatial position of an animal can be reliably decoded from the neuronal activity of several cell populations in the hippocampal formation [1–3]. For example, place cells in the hippocampus fire at only few locations in a spatial environment [4, 5] and the position of the animal can be readily read out from single active neurons. Grid cells of the medial entorhinal cortex (MEC) fire at multiple distinct places that are arranged on a hexagonal lattice [6, 7]. Although hexagonal patterns are abundant in nature and there exist well-studied physical theories for their emergence, the mechanistic origin of this neuronal grid pattern is still unclear. Initially it was suggested that they result from continuous attractor dynamics [8, 9] or superposition of plane wave inputs [10] and, based on circuit anatomy, place cells would then result from a superposition of many grid cells [11, 12]. More recent experiments, however, reported place cell activity without intact grid cells, such that grid cells are not the unique determinants of place field firing [13–17]. Conversely, it would thus be possible that grid fields may arise from place field input as suggested in [18–20]. The biological mechanisms proposed by these latter theories, however, remain hypothetical. In the present Letter, we propose a learning rule for grid cells based on the individual spike timings of place cells using spike-timing dependent synaptic plasticity (STDP) [21–23]. The theory thereby predicts that the observed temporal hippocampal firing patterns (phase precession and theta-scale correlations; see below) [24–26] translate the temporal proximity of sequential place field spikes into spatial neighborhood-relations observed in grid-field activity. For our model to work, we only have to assume that the synaptic plasticity rule averages over a sufficiently long time interval.

## Model

We use the classical formulation of pairwise additive STDP [22, 27], where the update of a synaptic weight *J_n_*, *n* = 1, …, *N* at time *t* is computed as [22]
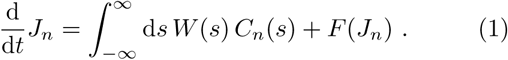

*C_n_*(*s*) denotes the time averaged correlation function between the spike train of presynaptic neuron *n* and the postsynaptic neuron, the learning window *W*(*s*) describes the update of the synaptic weight as a function of the time difference *s* between a pair of pre- and postsynaptic action potentials, and the function *F* implements soft bounds for the weight increase. The dynamics is further constrained such that weights cannot become negative.

To be able to treat eq. (1) analytically, we use a linear Poisson neuron model, i.e., the mean firing rate of the postsynaptic neuron *E*(*t*) = **J** • **H**(t) results from a weighted sum of hippocampal firing rates **H** = (*H*_1_(*t*), …, *H_N_*(*t*))^T^. Under these assumptions *C_n_*(*s*) can be approximated for large *N* [22] as 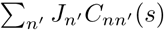 with
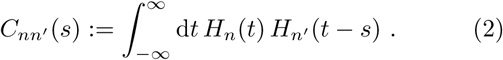

Inserting the correlation functions from eq. (2) into the weight dynamics from eq. (1) yields
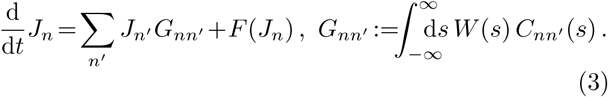

Following [28, 29] we introduce the quadratic stabilization term *F*(*J*) = *F*_o_ *J* (*K – J*), *F*_o_ > 0 that implements a soft upper bound.

As an input to the postsynaptic neuron, we consider a population of *N* hippocampal place cells. The firing of these neurons is characterized by a bell shaped envelope modulating the spatial path 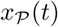 and oscillations in time *t* (Fig. 1A),
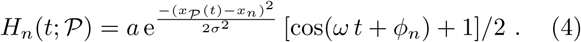

**FIG. 1.**
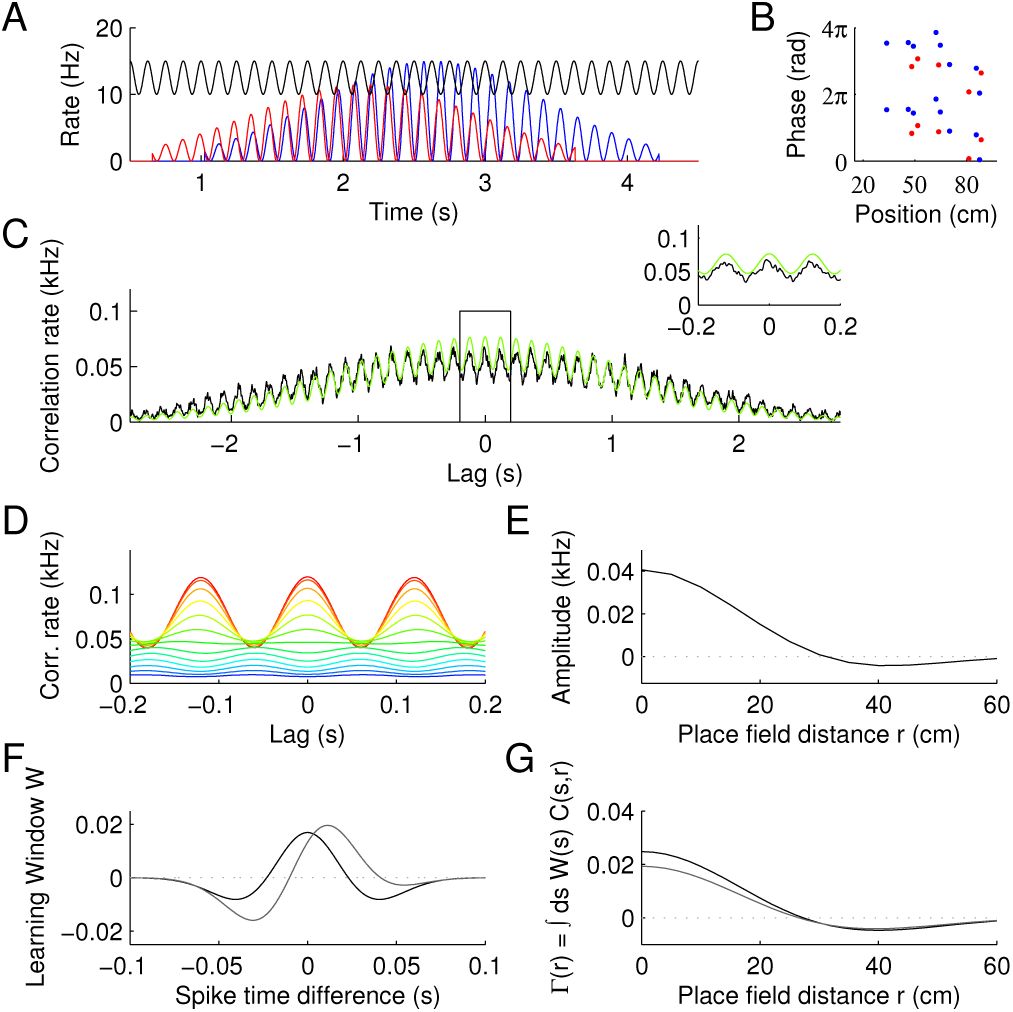
(A) Poisson model of place field firing for two place cells (red, blue), and slower theta oscillation (black). (B) Phase precession resulting from the path used in A. (C) Correlation of spike trains for the cells in A averaged over 2-d trajectories (random walk) (black), and from eq. (5) (green). Inset magnifies lag 0. (D) Correlation functions for different place field distances (red: autocorrelation; blue: 60 cm). (E) Correlation amplitude as function of place field distance. (F) Examples of STDP learning windows; see eq. (9) and below. (G) Γ kernels for correlation function from C-E and learning windows from F.

The oscillation frequency *ω* of a neuron is slightly higher than the frequency *ω_θ_* of the theta oscillation (~ 8 Hz) in the local field potential giving rise to a phenomenon called theta phase precession (Fig. 1B): spikes early in the field come at later phases than spikes late in the field [24]. During traversal of a place field, phase precession spans a whole theta cycle [30]. Thus the two frequencies have to relate to each other like 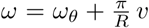, with *R* denoting the distance from the place field center at which the firing rate has decreased to 10% (i.e., 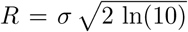), and *v* denoting the running speed, which we fix at 25 cm/s. At each individual entry into the place field, the phase of the cellular oscillation is reset to phase zero with respect to the theta oscillation phase *ϕ_θ_* by fixing *ϕ_n_* = (*ϕ_θ_* − 2*π*) *ω*/*ω_θ_.*

To obtain a closed expression for the correlation function *C_nn_*_′_(*s*) (Fig. 1C), the time average in eq. (2) is performed over all straight paths 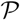 crossing the center of the field overlap. For place fields with identical width *R*, firing rates *a*, and at small lags *s*, we obtain
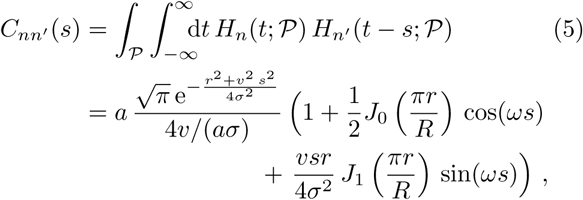

with place field distance *r* = |*x_n_*−*x_n_*_′_|, and *J*_0,1_ denoting Bessel functions of first kind (see Supplementary Material at [URL] for derivation).

In contrast to 1-d fields where the distance of place fields is reflected by the lag of the correlation peak (theta compression) [25], correlation functions in 2-d are symmetric because of the symmetry of the path, however, the distance of the place field centers is encoded in the amplitude of the correlation peak at lag 0 (Fig. 1D,E).

## Weight Dynamics

Assuming that the putative grid cells receive inputs from a large number *N* ≫ 1 of place cells that sufficiently cover the encoded area, we replace the presynaptic index *n* by the position of the place field center *x*, i.e., *G_nn_*_′_ → Γ(|*x_n_*−*x_n_*_′_|), and thereby translate the learning eq. (3) to continuous coordinates,
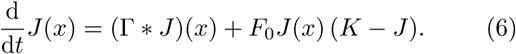

Examples of the convolution kernel Γ(|*x*|) for different learning window functions *W* are depicted in Fig. 1F, G. The development of the weights follows the pattern formation principles of a lateral inhibition system [31]. Indeed, the integro-differential equation (IDE) (6) involves non-local interactions effectively implemented through the convolution kernel, inducing a strong close range potentiation and a weaker long range depression of neighbouring synapses, as observed in the typical shape presented in Fig. 1G. A general window-dependent kernel can be obtained for the correlation function from eq. (5) as
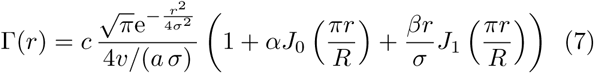

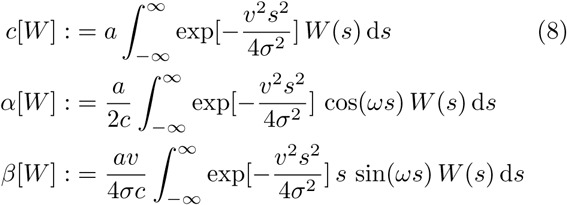

which can take a Mexican-hat type shape for qualitatively different learning window functions *W*, due to the symmetry of the correlation function (Fig. 1F, G). To see this, we can regard Hebbian-like windows to be modelled as the product of a Gaussian and a polynomial of some order *m*, *W*(*s*) = exp[−*s*^2^/(2*ρ*^2^*μ*^2^)] *P_m_*(*s/ρ*). The functionals defined in eq. (8) then inherit the symmetries from the cross-correlation, since all of the odd terms in the polynomial cancel out during integration. Thus, in subsequent numerical investigations we focus on windows up to second polynomial order
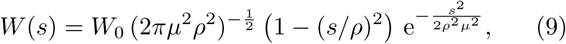

whose free parameters *ρ* and *μ* determine its zeroes *s*_0_ = ±*ρ* and negativity ∫*W* = *W*_0_(1−*μ*^2^). In Figure 1F,G we used *ρ* = 23 ms, *μ* = 1.025 and added a linear term *s*/*ρ* to the polynomial to get the asymmetric window (grey lines). The W-dependent functions *c*, *α*, and *β* defining the kernel Γ are given in the Supplementary Material [URL] .

A numerical evaluation of the learning IDE (6) with periodic boundary conditions reveals that the spatially isotropic kernel Γ can result in hexagonal packing structures (Fig. 2A, B). Simulations of spiking Poisson neurons confirm these predictions of the meanfield theory (Fig. 2A). As indicated by the kernel function in eq. (7), the grid spacing only depends on the spatial scale *σ* of the place fields in the input (Fig. 2C).

**FIG. 2.**
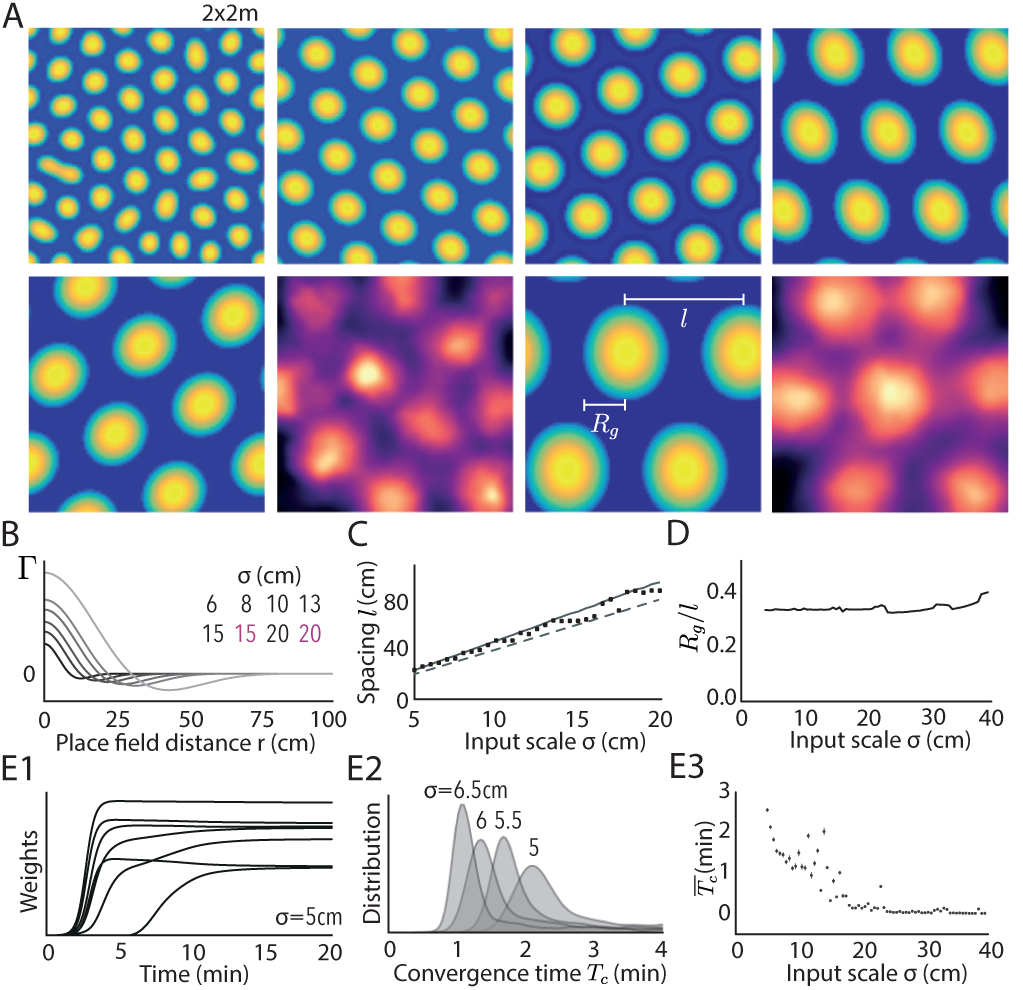
Weight patterns and input scale. (A) Asymptotically stable weight functions *J*(*x*) for place field widths *σ* as indicated in (B) together with the respective Γ kernels (*μ* = 1.025, *ρ* = 23 ms, *σ* indicated by grey level). Firing rates of spiking simulations are colored in red (for numerical details see Supplementary Material [URL]). (C) Grid spacing *l* (see A) scales linearly with place field scale *σ.* Dots correspond to weight patterns from numerical solutions. Dashed and solid lines indicate theoretical estimates 2π/*k_m_* and 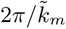, see *Linear theory at early times* and *Positivity constraint.* (D) Estimated ratio of field radius *R_g_* (see A) to grid spacing *l*, 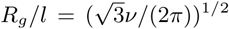 is independent of *σ.* (E_1_) Temporal evolution of randomly sampled weights. (E_2_) Distribution of weight convergence times *T_c_* for different input scales *σ* as indicated and the respective means (E_3_).

An experimentally accessible quantity to compare our model results to is the ratio of grid field radius *R_g_* to grid spacing *l* as indicated in Fig. 2A. In experiments, the ratio *R_g_*/*l* for grid cells has been determined to be about 0.3 [7], and for a perfectly hexagonal grid *R_g_*/*l* relates to the fraction *v* of field size per area as 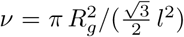.

The field fraction *v* can be readily accessed from our numerics as the fraction of non-zero synaptic weights, and for the used learning window fits the experimentally obtained *R_g_*/*l* (Fig. 2D) for all choices of *σ*. Finally, learning converges faster for large spacing (large *σ)* consistent with a larger amplitude of Γ [eq. (7) and Fig. 2B,E].

## Linear theory at early times

Some analytical understanding of the weight dynamics from eq. (6) can be gained from a neural field theory approach [31–35]. In this framework, we can neglect the effect of the nonlinearities at early times, and focus only on the convolution term Γ * *J*. The emerging dynamics can be readily understood by looking at the evolution of the weights in Fourier space 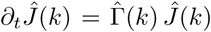, which makes evident that the wave number *k_m_* maximizing the kernel Fourier transform 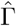 (see Supplementary Material [URL]) will exponentially overgrow all other modes (if 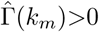), thus setting the initial periodicity of the pattern.

In a finite region of the parameter space (*μ, p*) of the learning window, the bimodal (Mexican-hat) shape of Γ ensures the existence of a Turing instability, i.e., a transition to single maximum of 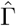 (with 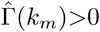) at a non-zero *k_m_*>0 (Fig. 3A-C). Similar to previous work on pattern formation in lateral inhibition systems (e.g. [36]), the permitted parameter region *(k_m_* exists and is positive) gives rise to stripe-like and hexagonal patterns (Fig. 3D). In the Supplementary Material [URL], we also provide a complementary description of the pattern formation process based on an approximation of eq. (6) by a partial differential equation (Swift-Hohenberg equation [37]), which corroborates the results from the linear theory.

**FIG. 3.**
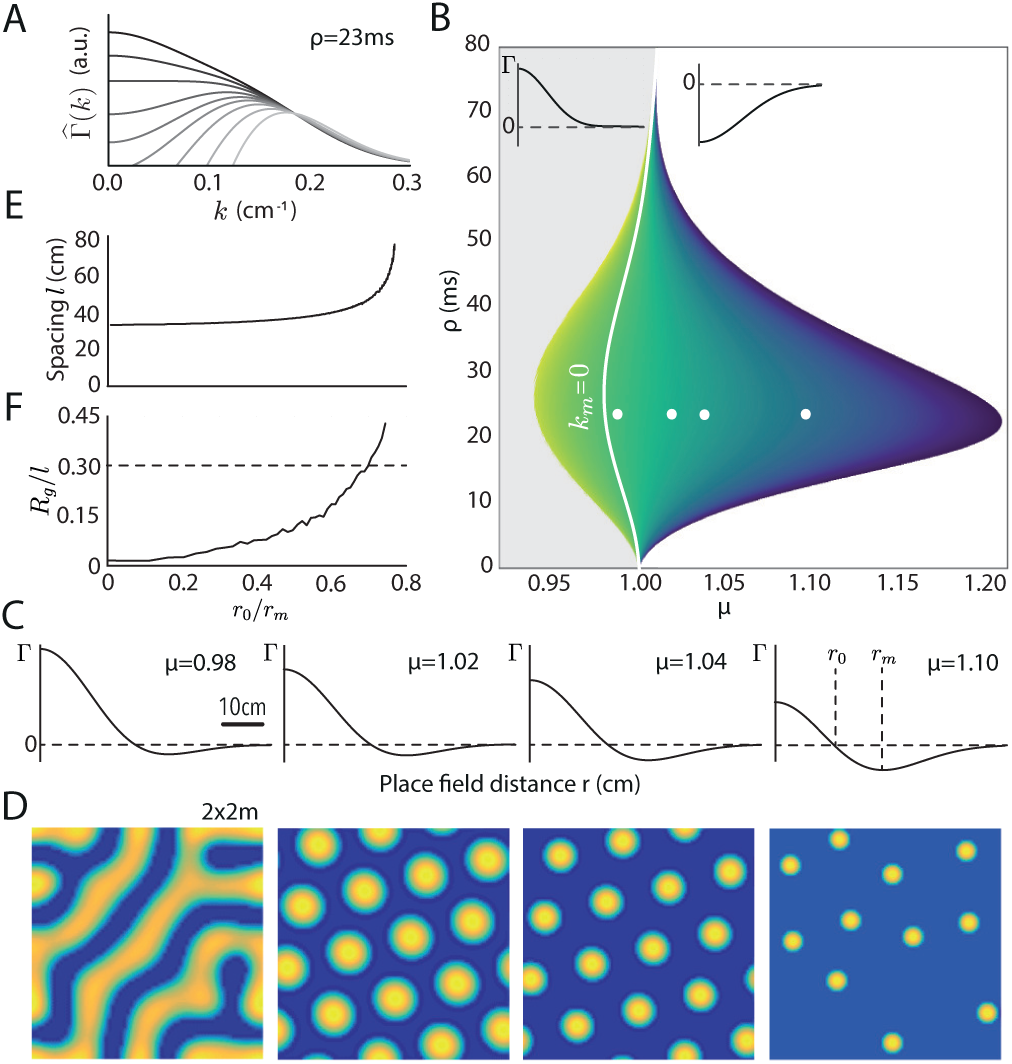
Permitted learning windows. (A) Fourier transforms of Γ for *μ* = 0.95 (black) to 1.20 (light grey). (B) Region of structure formation in (*μ*, *ρ*) space (for *σ* = 10 cm). Nontrivial patterns appear to the right of the white solid line (*k_m_* = 0), where the selected wavelength is positive. Colored region indicates bimodal Γ. The color encodes the shape factor *r*_0_/*r_m_* (0: blue; 1: yellow). Grey and white areas correspond to regions where Γ is all positive or all negative. (C) Four examples of Γ kernels corresponding to the white dots from B in the same order. (D) Weight patterns for kernels from C. (E) Predicted spacing 2π/*k_m_* as a function of *r*_0_/*r_m_* obtained for all combinations of *μ* and *ρ* from the region right of the solid line in A. (F) Estimated ratio of field size to grid spacing as a function of *r*_0_/*r_m_* (from simulated patterns). Dashed line indicates experimental value [7].

To connect the resulting patterns to other feed-forward models of grid field formation, we parameterize Γ by the shape factor *r*_0_/*r_m_* (Fig. 3C), which is the fraction between the zero and the minimum of Γ. The shape factor *r*_0_/*r_m_* reduces the two-parameter learning window to a single qualitatively descriptive parameter, which can be used to describe the bimodal kernel Γ independently of the hypothesized biological mechanism. If *r*_0_/*r_m_* is large (~ 0.8), Γ shows only little negativity and the emerging pattern is stripe-like (Fig. 3C, D), if *r*_0_/*r_m_* is small, Γ exhibits strong negativity, the firing fields become dispersed and the pattern looses hexagonality. Hexagonal patterns arise for *r*_0_/*r_m_* roughly between 0.65 and 0.75 (Fig. 3D). In this region, the shape factor virtually completely determines the geometrical properties of the steady state (Fig. 3E, F). Values of *r*_0_/*r_m_* that give rise to hexagonal grids can also be identified via the ratio of field width per grid spacing *R_g_*/*l.* According to our theory, the experimentally observed value 0.3 [7] is achieved with a shape factor of about *r*_0_/*r_m_* = 0.7 (Fig. 3F). For higher values of *r*_0_/*r_m_*, *R_g_*/*l* increases to a point where a periodic pattern cannot further dissociate into disjoint fields and the stable pattern becomes stripe-like. For lower *r*_0_/*r_m_*, *R_g_*/*l* decreases, and at some point, the small fields no longer repel each other strongly enough to produce a symmetrical arrangement.

## Positivity constraint

The grid spacing *l* predicted by the linear theory, however, consistently underestimates the spacing derived from the numerical solution of the mean field dynamics (Fig. 2C). The reason for this error is that, after the initial growth phase, the synaptic weights are influenced by the non-linearities, most importantly the constraint that they cannot become negative.

The impact of this positivity constraint can be intuitively understood if we interpret the convolution Γ * *J* as an operation that detects the best overlap of an oscillatory pattern *J* ∝ cos(*kx*) with a given kernel Γ. However, after the lowest weights reach zero they stop contributing to the convolution and a slightly lower wave number 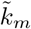 maximizing
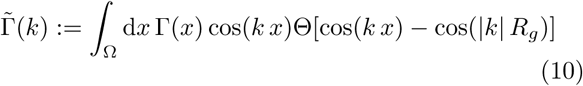

will be favored as the fastest growing mode (Θ denoting the Heaviside function). Similarly, a particular field size *R_g_* maximizing 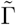 will be selected. In the experimentally relevant case |*k*| *R_g_* = 2π*R_g_*/*l* = 2π x 0.3, numerical maximization of eq. (10) yielded the predicted wave number 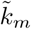 (solid line in Fig. 2C), which excellently agrees with the numerical solutions of the meanfield dynamics.

## Conclusion

For a large variety of STDP windows, the spike-timing correlations of 2-d place cells can account for a feed-forward learning of hexagonal grid patterns. Synaptic plasticity thereby averages over running trajectories of tens of minutes, hence, translating the temporal correlations into a dense code for space. Our model thus predicts that grid cells are generated in the output structures of the hippocampus, e.g., the deep layers of the medial entorhinal cortex [38] or the parasubiculum [39]. While our linear theory provides a good prediction of grid spacing as well as for conditions that permit structure formation, determining the boundary between hexagonal and stripe-like patterns is less straight-forward and has to take into account the non-linearities. The standard approach, non-linear bifurcation analysis [36, 40, 41], is difficult because of the strong non-linearity introduced via the positivity constraint, which strongly influences the selection of the final pattern. Despite this drawback, our model provides a universal framework in that it encompasses current models of grid field formation that can be mapped to convolutions with Mexican hat-type kernels that give rise to a Turing instability.

The authors are grateful to Andreas Herz and Anton Sirota for discussions and Martin Stemmler for comments on the manuscript. This work was funded by the German Research Association (DFG), Grant No. LE2250/5-1.

